# α-Synuclein polymorphism determines oligodendroglial dysfunction

**DOI:** 10.1101/2021.07.09.451731

**Authors:** Benedikt Frieg, James A. Geraets, Timo Strohäker, Christian Dienemann, Panagiota Mavroeidi, Byung Chul Jung, Woojin S. Kim, Seung-Jae Lee, Maria Xilouri, Markus Zweckstetter, Gunnar F. Schröder

## Abstract

Synucleinopathies, such as Parkinson’s disease (PD) and Multiple System Atrophy (MSA) are progressive and unremitting neurological diseases. For both PD and MSA, α-synuclein fibril inclusions inside brain cells are neuropathological hallmarks. In addition, amplification of α-synuclein fibrils from body fluids is a potential biomarker distinguishing PD from MSA. However, little is known about the structure of α-synuclein fibrils amplified from human samples and its connection to α-synuclein fibril structure in the human brain. Here we amplified α-synuclein fibrils from PD and MSA brain tissue, characterized its seeding potential in oligodendroglia, and determined the 3D structures by cryo-electron microscopy. We show that the α-synuclein fibrils from a MSA patient are more potent in recruiting the endogenous α-synuclein and evoking a redistribution of TPPP/p25α protein in mouse primary oligodendroglial cultures compared to those amplified from a PD patient. Cryo-electron microscopy shows that the PD- and MSA-amplified α-synuclein fibrils share a similar protofilament fold but differ in their inter-protofilament interface. The structures of the brain-tissue amplified α-synuclein fibrils are also similar to other *in vitro* and *ex vivo* α-synuclein fibrils. Together with published data, our results suggest that αSyn fibrils differ between PD and MSA in their quaternary arrangement and could further vary between different forms of PD and MSA.

## Introduction

α-synucleinopathies are neurodegenerative diseases that feature the misfolding and abnormal aggregation of the presynaptic protein α-synuclein (αSyn) into megadalton-size inclusion bodies within the brain. Though sharing a similar molecular mechanism of seeding and assembly, the etiology and phenotype of the α-synucleinopathies are different, as are the roles of αSyn as both an effector of neurotoxicity and as a mediator of the pathogenicity and disease progression. These differences are manifest in macroscopic structural differences in the deposited aggregates (Peng et al., 2018; Strohäker et al., 2019), ascribed to different conformational polymorphs of self-propagating αSyn (Strohäker et al., 2019). *In vitro*-amplified polymorphs have been shown to differ in their seeding and self-propagation behavior *in vivo*, inducing polymorph-specific pathology and neurotoxic phenotypes (De Giorgi et al., 2020; Holec & Woerman, 2020; Lau et al., 2020; Lempriere, 2020; Strohäker et al., 2019; Suzuki et al., 2020; Van der Perren et al., 2020)

The hallmarks of Parkinson’s disease (PD) and dementia with Lewy bodies (DLB) are the presence of Lewy bodies within neurons, or Lewy neurites, filled with long αSyn filaments. For multiple system atrophy (MSA), αSyn is predominantly deposited in inclusions within oligodendrocytes, with limited pathology in neurons. αSyn also is the etiological agent of MSA and has been shown to directly transmit and self-propagate misfolding when transferred to transgenic mice or cell cultures. However, for the Lewy body diseases PD and DLB, αSyn has not been shown to transmit disease *in vitro* and *in vivo* directly (Recasens, Ulusoy, Kahle, Di Monte, & Dehay, 2018; Woerman et al., 2015). Besides the spreading of amyloid fibrils, smaller αSyn oligomers play an important role in the pathogenesis and progression of PD, DLB, and MSA (Chiti & Dobson, 2017; Roberts & Brown, 2015).

Monomeric αSyn is an intrinsically disordered, highly flexible protein in solution that exchanges between many different conformations. Based on its amino acid composition, three distinct regions can be distinguished: a structurally heterogeneous amphipathic N-terminal domain (aa 1–60), a more hydrophobic central core region (aa 61–95), and an acidic C-terminus (aa 96–140) (Der-Sarkissian, Jao, Chen, & Langen, 2003). Much of the N-terminal and central regions consist of seven imperfect repeats, with the consensus sequence KTK(E/Q)GV. These repeats form a lipid-binding N-terminal helix (Davidson, Jonas, Clayton, & George, 1998). The hydrophobic non-β-amyloid component neighbors this in the central region (aa 61–95), of which the inclusion of 12aa is both necessary and sufficient for fibrils to form (aa 71–82).

Several αSyn fibril polymorph structures have been determined. Notably, there are structural differences between the *in vitro* aggregated fibrils and the fibrils extracted from patient tissue, even if *ex vivo* seeds are used for fibril propagation, which affects the extended folds and asymmetric packing of the protofilaments (Lövestam et al., 2021). Besides calling into question the continued use of non-brain-amplified αSyn to study PD and MSA, there is the broader consideration that additional factors are underpinning the process of fibrilization *in vivo*. In addition, these structural studies have suggested that a consensus protofilament fold, which consists of a common protofilament core but with different inter-protofilament interfaces and terminal arms, might be conserved across αSyn fibril polymorphs derived from different sources (Lövestam et al., 2021; Schweighauser et al., 2020). This raises the question of what factors mediate the assembly of the common αSyn protofilament core *in vivo* and control their overall arrangement within the fibril. Posttranslational modifications such as phosphorylation, ubiquitination, and nitration are seen in filamentous αSyn *in vivo*, and have been seen to play roles in the propagation of misfolding and aggregation (Sorrentino and Giasson, 2020), as does the proteolytic truncation of αSyn, particularly of the C-terminus (Games et al., 2014). Similarly, alternative splicing of the *SNCA* gene encoding for αSyn gives rise to four isoforms, which all have different propensities for aggregation (Beyer & Ariza, 2013). Synucleopathies manifest with pathognomonic assemblies in distinct cell types, where cellular factors differ; the physiological expression of αSyn varies in different brain regions and other cell types (Courte et al., 2020; Taguchi, Watanabe, Tsujimura, & Tanaka, 2016). In particular, differences between the cellular milieus of oligodendrocytes compared with other brain cells may be crucial for MSA pathogenesis and progression. Hence, specific factors within oligodendrocytes might play a role in imparting an MSA-amplified conformation on αSyn fibrils and inducing their characteristic direct self-propagation (Peng et al., 2018).

In addition to the putative structural differences attributed to different α-synucleinopathies it also appears that αSyn can adopt a range of structural polymorphs within a single disease (Schweighauser et al., 2020), though this might vary between clinical cases. Interestingly there could be more heterogeneity among PD than MSA fibrils, which may be linked to the greater variety of disease phenotypes in PD (Klingstedt et al., 2019; Strohäker et al., 2019). Differences in the structure of αSyn fibrils could account for the differing pathogenesis and disease progression by being a result of different post-transformational modifications, in particular phosphorylation and ubiquitination (Suzuki et al., 2020), where for example, it has been seen that resulting differences in the C-terminus led to differential inhibition of proteasome activity.

Amplification of protein aggregates by protein misfolding cyclic amplification (PMCA) and RT-QuiC can be used to detect αSyn aggregates in cerebrospinal fluid (Fairfoul et al., 2016; Kang et al., 2019; Shahnawaz et al., 2017) and tissue lysates (Becker et al., 2018; Jung et al., 2017; Sano et al., 2018) with high sensitivity and specificity. Remarkably, work by Shahnawaz *et al*. suggests that PD and MSA might be distinguished using αSyn from the cerebrospinal fluid as seed and recombinant αSyn as the substrate for protein aggregate amplification (Shahnawaz et al., 2020).

To gain insight into the molecular structure of αSyn aggregates associated with PD and MSA, we amplified αSyn fibrils from brain extracts of patients pathologically confirmed with PD and MSA, evaluated their seeding potential in oligodendroglia and solved their cryo-EM structures. The structures of PD- and MSA-amplified αSyn fibrils reveal several similarities and dissimilarities, allowing discrimination between PD- and MSA-amplified αSyn fibrils. Notably, seeded recombinant αSyn fibrils partially replicate the structure of fibrils found in MSA patients. Still, the influence of additional molecules or posttranslational modifications of αSyn may be required for the entire replication.

## Results

### MSA-amplified *α*Syn fibrils are more active in oligodendroglia than PD-amplified fibrils

We seeded fibril formation of recombinant αSyn through the addition of PMCA-products, which were previously generated from the homogenized brain tissue of a PD and a MSA patient (PD and MSA patient #1 in (Strohäker et al., 2019)). Hydrogen-deuterium exchange coupled to NMR spectroscopy as well as fluorescence dye binding had shown that the brain-tissue amplified αSyn fibrils (further termed PD- and MSA-amplified αSyn fibrils) differ in their structural properties (Strohäker et al., 2019).

To gain insight into potential differences in the cellular activity of these PD- and MSA-amplified αSyn fibrils, we added each fibril sample to differentiated murine primary oligodendroglial cultures. The distinct profiles of PD- and MSA-amplified fibril strains were further validated by the differential pathology-related responses observed in these cultures upon their inoculation with the patient amplified fibrils. In particular, our results demonstrate that MSA-amplified fibrils display higher potency in seeding the endogenous oligodendroglial αSyn and promoting the redistribution of the oligodendroglial-specific phosphoprotein TPPP/p25α from the myelin sheath to the cell soma, as compared to PD-amplified fibrils (**Figure 1**). Both events are considered to play an important role in the cascade of events leading to oligodendroglial dysfunction and neuronal demise underlying MSA pathology.

**Figure 1:**
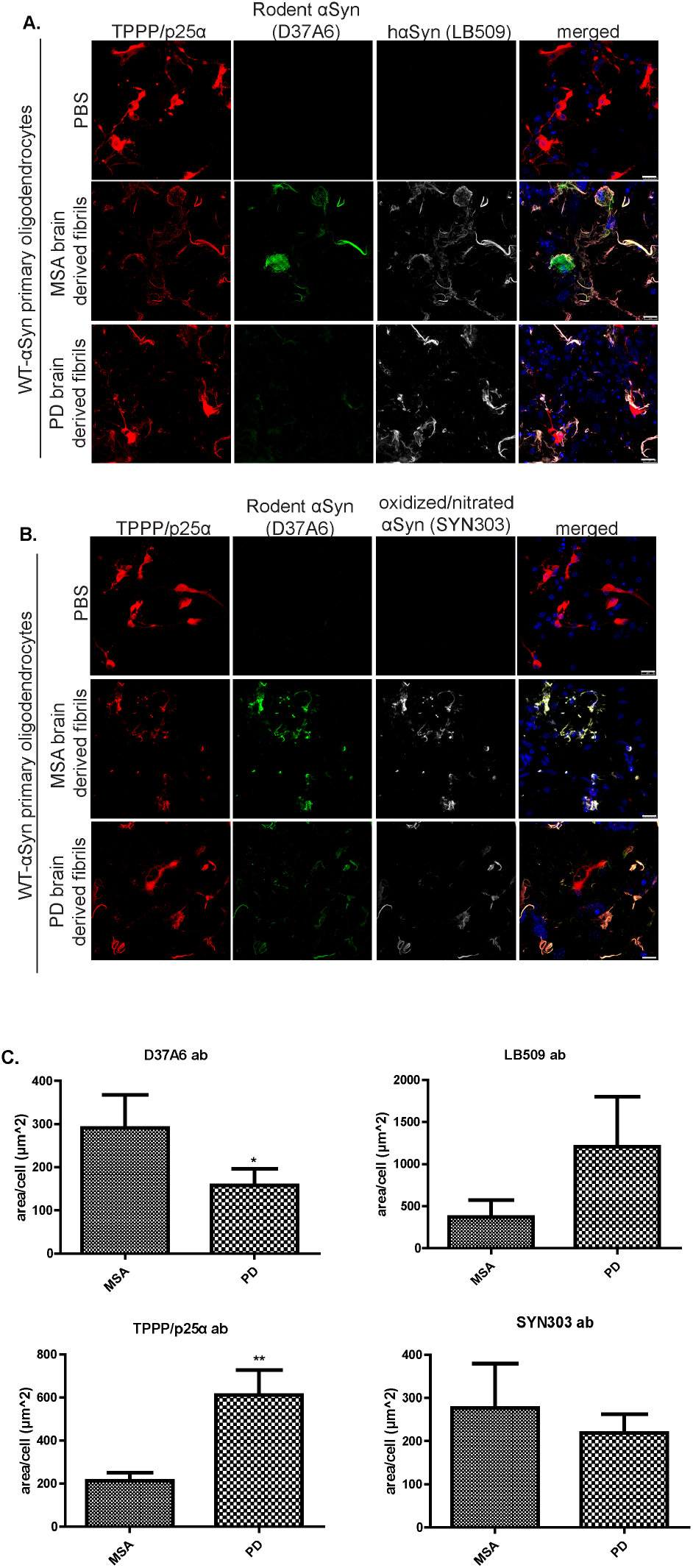
Human *α*Syn fibrils amplified from a MSA patient are more potent in recruiting the endogenous oligodendroglial *α*Syn and evoking a redistribution of TPPP/p25*α* protein in mouse primary oligodendroglial cultures, compared to those amplified from a PD patient. Both MSA- and PD-brain derived fibrils lead to the formation of pathological αSyn species (oxidized/nitrated αSyn, SYN303 ab). **(A-B)** Representative immunofluorescence images of mouse primary oligodendrocytes treated with 0.5 µg human MSA or PD fibril strains for 48 h (or PBS as control) using antibodies against TPPP/p25α (red), endogenous rodent αSyn (D37A6 antibody, green), human αSyn (LB509 antibody, gray in A), oxidized/nitrated αSyn (SYN303 antibody, gray in B) and DAPI (shown in blue) staining as nuclear marker. Scale bar: 25 µm. **(C)** Quantification of the endogenous rodent αSyn (upper left), human αSyn (upper right), TPPP/p25α (lower left) and oxidized/nitrated αSyn (lower right) protein levels in mouse primary oligodendrocytes, measured as µm^2^ area surface/cell following their treatment with 0.5 µg MSA- or PD-brain derived fibrils for 48 h. Data are expressed as the mean ± SE of three independent experiments with duplicate samples/condition within each experiment; *p<0.05; **p< 0.01, by Student’s unpaired *t* test.

### αSyn fibrils reveal a common protofilament fold but a distinct inter-protofilament interface

To elucidate why primary oligodendroglia respond differently to PD- and MSA-amplified fibrils, we used cryo-EM and solved the 3D structure of the same samples of brain-tissue amplified αSyn fibrils, which showed the differential response in the oligodendroglia. Visual inspection of the micrographs revealed the presence of a single main fibril in both cases, with measured crossover distances of ∼1000 Å and ∼1200 Å for PD- and MSA-amplified αSyn fibrils, respectively (**Figure 2A, B, Figure S1**), suggesting for two distinct fibril polymorphs specific for either disease. As to MSA-amplified fibrils, we also found non-twisted fibrils (**Figure S1**), which were considered preparation artifacts from interactions with the air-water interface, as suggested previously (Lövestam et al., 2021).

**Figure 2:**
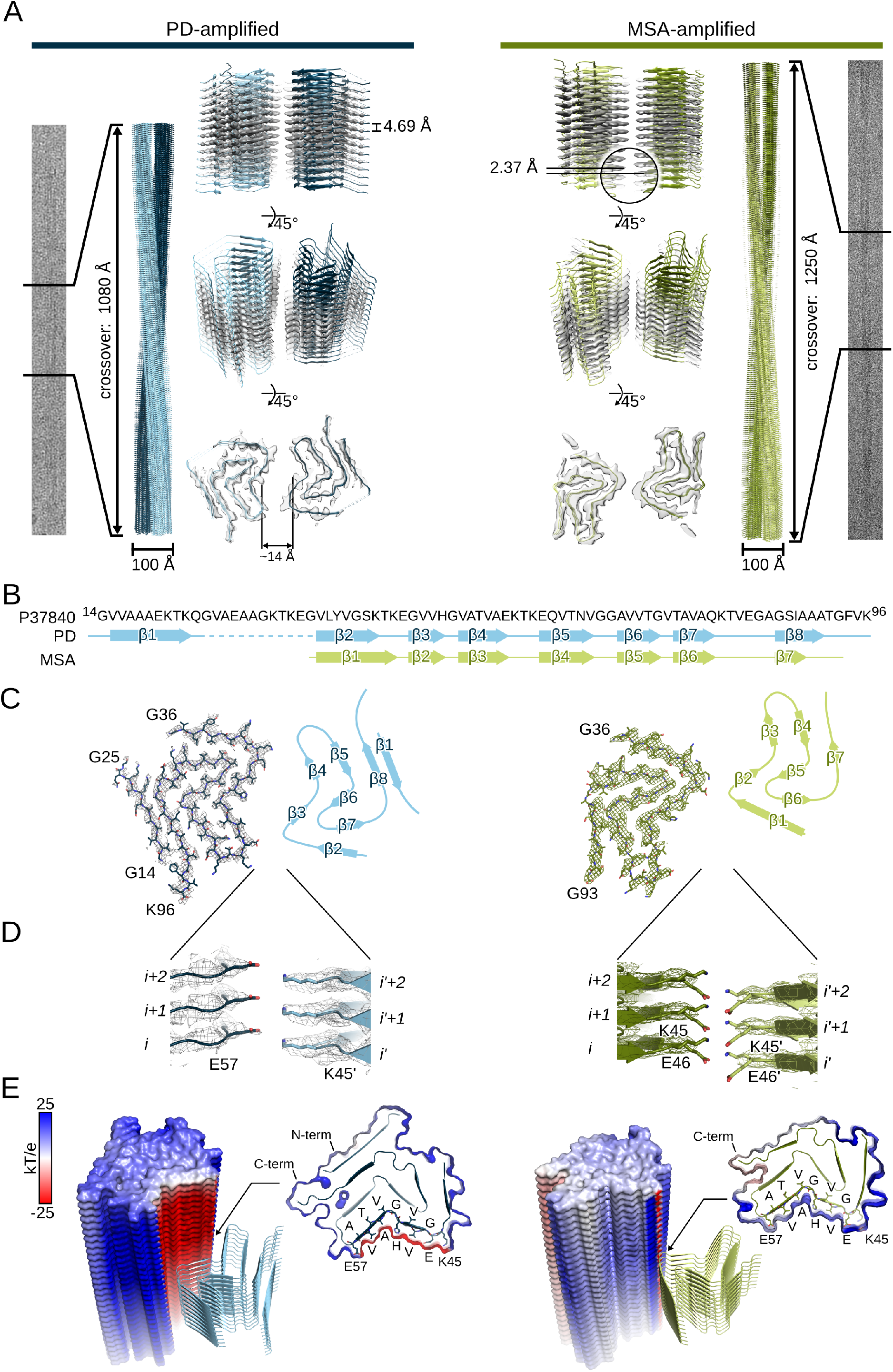
Cryo-EM structures of amplified αSyn fibrils. **A**: The cryo-EM structures of amplified αSyn fibrils from PD (left) and MSA (right) brain. From the outside to the inside, the panels show extracts from a representative micrographs with amplified fibrils, a full cross over (180° turn) of the reconstructed fibril, with two protofilaments colored in different shades of blue (in the case of PD) or green (in the case of MSA), and semitransparent surfaces overlaid with their atomic models viewed from different angles. As to the MSA fibril, both protofilaments arrange in an approximate 2_1_ screw symmetry. **B**: Amino acid sequence of αSyn (from G14 to K96; based on UniProt: P37840) with a schematic depiction of the secondary structure of the protofilament fold. β-strands are shown as arrows and numbered from β1 to β8 (for PD) or β7 (for MSA), respectively. The region from V26 to E35 was not resolved (indicated by a dashed line). **C**: Top view onto two opposite subunits of the reconstructed PD (left; colored in shades of blue) and MSA (right; colored in shades of green). One protofilament is shown as mesh-stick representation, the other schematically depicted by its secondary structure matching the assignment in **B. D:** Close up view of the protofilament interface, with interface amino acids shown as stick models. **E:** The calculated electrostatic potential was mapped onto the surface of one protofilament and colored according to the color scale on the left. The central subunits of the other protofilament are shown as cartoon-model. Cross sections are shown as surface-cartoon model with amino acids forming the central negatively charged cavity in PD-amplified αSyn (from K45 to E57) labeled explicitly.

The 3D structures of PD and MSA-amplified fibrils were determined to a resolution of 3.3 Å and 3.0 Å, respectively, based on the gold-standard Fourier shell correlation 0.143 criterion (**Table 1, Figure 2**, and **Figure S2)**. In the case of the MSA-amplified fibril, however, the local resolution estimation revealed a high heterogeneity with a resolution of ∼2.9 Å in the inter-protofilament interface and >4.0 Å at the periphery of the fibril (**Figure S3**). Indeed, the side-chain densities at the periphery of the fibril are not well resolved. For the PD-amplified fibril, by contrast, the local resolution estimates are more homogenous and, hence, the reconstructed map revealed clear side-chain densities. We, thus, assume that the overall fold of MSA-amplified fibril tends to be more flexible compared to the PD-amplified fibril. Still, both reconstructed maps show a clear β-strand separation along the helical axis and reveal that two intertwined protofilaments form both types of fibrils (**Figure 2A, B**). For PD-amplified fibrils, the protofilaments are related by C2 symmetry with a helical rise of 4.68 Å and, assuming left-twisting handedness, a helical twist of -0.78°. Interestingly, both protofilaments form a cavity of ∼14 Å in diameter (**Figure 2A**). By contrast, in MSA-amplified fibrils, the protofilaments are related by an approximate 2_1_ screw symmetry with a helical rise of 2.37 Å and twist of 179.66°, again assuming left-twisting handedness. Thus, the refined crossover distances are in excellent agreement with the measurements from the micrographs. In both cases, the fibril width is ∼100 Å. Due to the lack of twist, we were not successful at solving the 3D structure of the non-twisted MSA-amplified fibrils (Lövestam et al., 2021; Schweighauser et al., 2020). Local resolution estimation revealed a resolution of ∼2.9 Å in the inter-protofilament interface and ∼4.0 Å at the periphery of the fibril (**Figure S3**). Hence, the reconstructed maps also show apparent side-chain densities. Interestingly, both amplified αSyn fibrils reveal several similarities and dissimilarities, allowing clear distinction between both types of fibrils.

**Table 1.** Cryo-EM structure determination statistics.

| | PD-amplified αSyn | MSA-amplified αSyn |
| --- | --- | --- |
| **Data collection** |  |  |
| Microscope | Titan Krios G2 | Titan Krios G2 |
| Voltage [keV] | 300 | 300 |
| Detector | K3 | K3 |
| Magnification | 81,000 | 81,000 |
| Pixel size [$\text{\AA}$] | 1.05 | 1.05 |
| Defocus range [$\mu\text{m}$] | -0.7 to -2.9 | -0.7 to -2.9 |
| Exposure time [s/frame] | 2.2 | 2.2 |
| Number of frames | 40 | 40 |
| Total dose [$\text{e}^-/\text{\AA}^2$] | 42.74<br>(1.07 $\text{e}^-/\text{\AA}^2/\text{frame}$) | 43.05<br>(1.08 $\text{e}^-/\text{\AA}^2/\text{frame}$) |
| **Reconstruction** |  |  |
| Micrographs | 6,780 | 4,474 |
| Box width [pixels] | 200 (1.05 $\text{\AA}/\text{pixel}$) | 500 (2.10 $\text{\AA}/\text{pixel}$)<br>250 (1.05 $\text{\AA}/\text{pixel}$) |
| Inter-box distance [pixels] | 18 | 50 |
| crYOLO<sup>a</sup> picked segments (no.) | 2,223,091 | 1,848,677 |
| Final segments [no.] | 23,937 | 25,692 |
| Final resolution [$\text{\AA}$]<br>(FSC=0.143) | 3.30 | 3.02 |
| Applied map sharpening<br>B-factor [$\text{\AA}^2$] | -112.5 | -101.8 |
| Symmetry imposed | C2 | C1 |
| Helical rise [$\text{\AA}$] | 4.68 | 2.37 |
| Helical twist [$^\circ$] | -0.78 | 179.66 |

<sup>a</sup> Ref. (Wagner et al., 2019)

**Table 2.** Model building statistics.

| | PD-amplified αSyn | MSA-amplified αSyn |
| --- | --- | --- |
| **Starting model [PDB code]** | 6SSX<sup>a</sup> | 6SST<sup>a</sup> |
| **Model composition** |  |  |
| Non-hydrogen atoms | 497 | 389 |
| Protein residues | 73 | 58 |
| **RMS deviations** |  |  |
| Bond lengths [Å] | < 0.05 | < 0.05 |
| Bond angles [°] | 2.01 | 1.68 |
| **Validation** |  |  |
| MolProbity score | 2.25 | 2.02 |
| Clashscore | 17.17 | 14.41 |
| Poor rotamers [%] | 0.00 | 0.00 |
| **Ramachandran plot** |  |  |
| Favored [%] | 91.30 | 94.64 |
| Allowed [%] | 8.70 | 5.36 |
| Disallowed [%] | 0.00 | 0.00 |
| **Model deposition** |  |  |
| PDB code | 7OZG | 7OZH |
| EMD code | 13123 | 13124 |

<sup>a</sup> Ref. (Guerrero-Ferreira et al., 2019)

PD- and MSA-amplified αSyn fibrils have a highly similar protofilament fold, with differences at their N-terminus and inter-protofilament interface. The PD-amplified protofilament extends from G14 to K96 and is composed of eight β-sheets, from which β2 to β8 are connected by a continuous backbone chain and β2/β3, β4/β5, and β6/β7 form a triple-stacked *L*-shaped core (**Figure 3C, D**). No apparent densities were found for the N-terminus from M1 to E13, the region from V26 to E35, and the C-terminus beyond K96, suggesting that these regions are more flexible. Two salt-bridges between K45 and E57’ and between E57 and K45’, harbored on two opposite subunits *i* and *i’* form the inter-protofilament interface (**Figure 3C**). The MSA-amplified protofilament extends from G36 to G93 and is composed of seven β-sheets, creating a continuous backbone chain and, similar to PD-amplified fibrils, the β-sheets form the same *L*-shaped core (**Figure 3A, B**). In contrast to PD-amplified fibrils, no backbone densities were found for the first 38 amino acids, similar to other structures of αSyn lacking the same region (Boyer et al., 2020). The inter-protofilament interface is formed by a sophisticated salt-bridge network, in which K45 from subunit *i* interacts with E46 on subunit *i+1* and E46’ on subunit *i’+1* (**Figure 3C**). Additionally, E46 on subunit *i* interacts with K45’ from subunit *i’*. The intact integrity of the network explains why the subunits are arranged in a staggered manner, adopting an approximate 2_1_ screw symmetry. Hence, PD- and MSA-amplified αSyn fibrils share the same protofilament fold, but the inter-protofilament interface and helical arrangement are different, which, in turn, determine additional structural properties.

**Figure 3:**
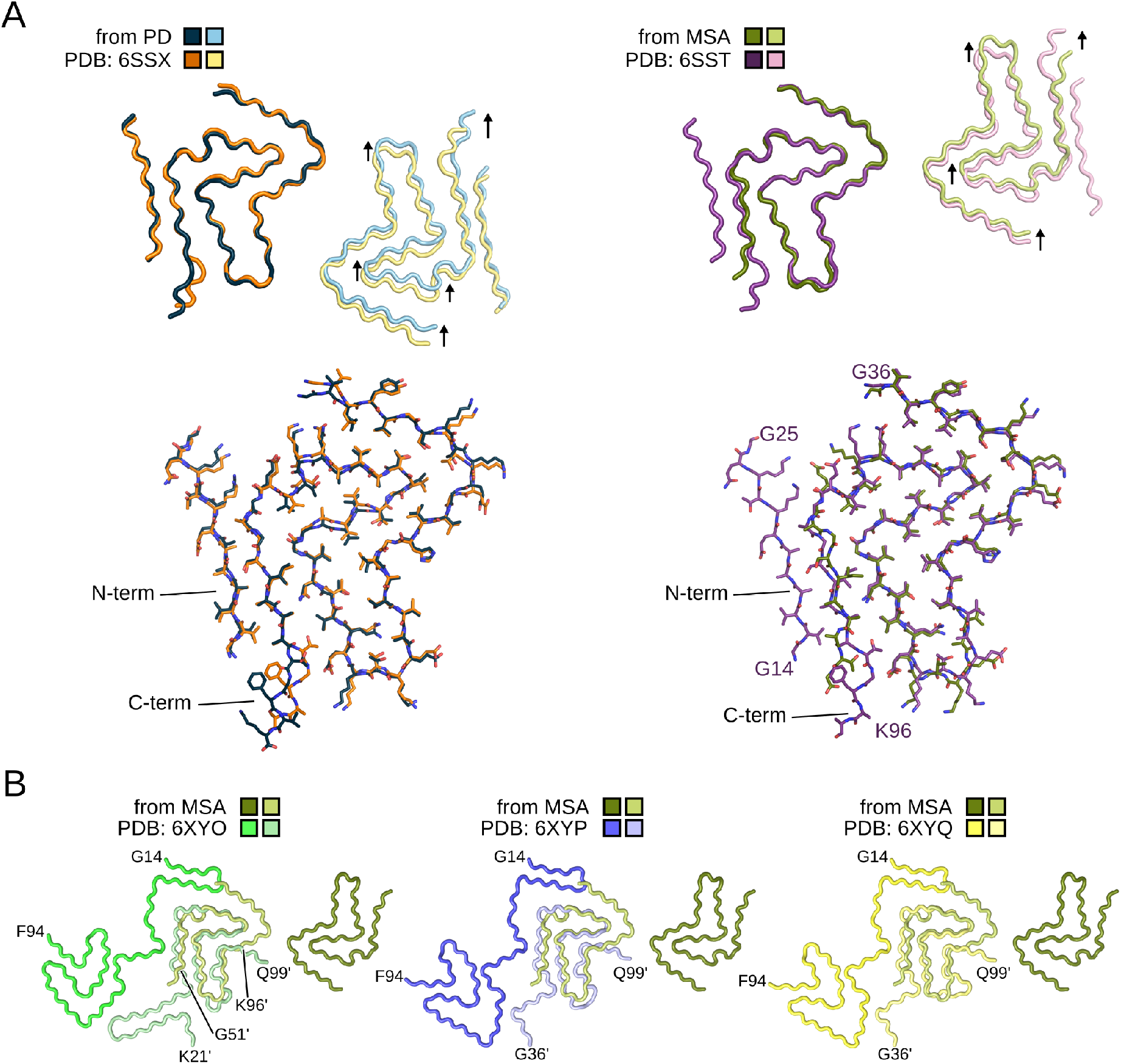
Amplified *α*Syn fibrils reveal a common protofilament fold. **A:** Overlay of amplified αSyn structures onto two opposite subunits extracted from recombinant *in vitro* aggregated αSyn fibrils with PDB IDs 6SSX and 6SST (Guerrero-Ferreira et al., 2019). The arrows indicated the relative shift between *in vitro* αSyn fibrils and the amplified αSyn fibrils (colored in light colores) after superimposing the Cα atoms of the opposite subunits (colored in dark colors). **B:** Overlay of amplified αSyn structures onto αSyn structures extracted from the brains of individuals with MSA with PDB IDs 6XYO, 6XYP, and 6XYQ (Schweighauser et al., 2020). The protofilaments are colored in different shades of green, blue, and yellow, with amino acids from subunit B labeled with an additional prime.

For both types of fibrils, we found pronounced discrepancies in the electrostatic surface potential. The PD-amplified fibril is markedly positively charged on the exterior surface, but the inter-protofilament cavity, interestingly, is oppositely charged (**Figure 3D, Figure S4**). The latter is impressive as the amino acids forming the cavity are, except from E46 and E57, not negatively charged. The same region is positively charged in the MSA-amplified fibril. Hence, the global distribution of the surface charges and the negatively charged cavity, in particular, likely results from the discriminative inter-protofilament interface. Although the exterior surface of the MSA-amplified fibril is also positively charged, the magnitude is reduced, particularly at the now almost neutral C-terminus, which the absence of the N-terminus may explain. Taken all together, PD- and MSA-amplified αSyn fibrils share the same protofilament fold, but the inter-protofilament interface and helical arrangement are different. The structural differences lead to pronounced differences in the fibril electrostatic interactions, which, in turn, may evoke the observed aberrant cellular responses associated with oligodendroglia dysfunction.

### Amplified *α*Syn fibrils reveal similarities to *in vitro* aggregated *α*Syn

The structure of our PD and MSA-amplified fibrils are highly similar to those reported for *in vitro* aggregated αSyn. We compared our αSyn structures to previously solved structures and used the Cα root mean square deviation (Cα RMSD) to measure structural similarity. Our PD-amplified αSyn structure is almost identical to the structure identified for *in vitro* aggregated recombinant wild-type αSyn, previously referred to as polymorph 2A (Guerrero-Ferreira et al., 2019) (**Figure 3A**). Cα RMSDs are 1.03 Å and 1.33 Å, considering one or both protofilaments, respectively (**Table S1**), suggesting only minor structural deviations at the C-terminus from T92 to K96 (**Figure 3A**). Although the protofilament fold and inter-protofilament interface are almost identical, the helical rise and twist are different (**Table S1**), resulting in a lateral displacement of the protofilaments relative to each other (**Figure 3A**). We assume that variations in the preparation protocols, e.g., seeded versus *de novo* assembly, denote a credible source for this discrepancy. However, the electrostatic surface potential of polymorph 2A is indistinguishable from our PD-amplified αSyn (**Figure S4**), corroborating our assumption that the positively charged exterior surface and the negatively charged inter-protofilament interface is the result of the helical arrangement.

Our MSA-amplified αSyn structure is similar to the structure identified for *in vitro* aggregated recombinant wild-type αSyn, previously referred to as polymorph 2B (Guerrero-Ferreira et al., 2019) (**Figure 3A**). Cα RMSDs are 1.47 Å and 1.74 Å, considering the resolved amino acids G36 to G93 of one or both protofilaments, respectively (**Table S2**), suggesting minor structural deviations. In contrast to polymorph 2B, the N-terminal region from G14 to G25 is not visible in the MSA-amplified structure, suggesting that the N-terminus tends to be more flexible. Although the protofilament fold and inter-protofilament interface are identical, helical rise and twist are different (**Table S2**), similar to what we found in the PD-amplified fibrils. The stabilized N-terminus in polymorph 2B, however, enhances the electrostatic surface potential compared to our MSA-amplified fibril (**Figure S4**). Interestingly, the charge distribution on the exterior surfaces is similar for polymorphs 2A and 2B (**Figure S4**), suggesting that the fold of the protofilament, which is identical in polymorphs 2A and 2B, determines the charge distribution on exterior surfaces.

Both of our amplified αSyn structures are also very similar to αSyn structures amplified from brain extracts of MSA patients (Lövestam et al., 2021) (**Table S1, S2**). Interestingly, superimposing one protofilament strand yields Cα RMSDs < 2 Å, but superimposing two opposite protofilament strands yields Cα RMSDs > 10 Å, suggesting that the global symmetry, rise, twist, and, thus, helical organization are different (**Table S1, S2**). Hence, our PD- and MSA-amplified structures show a similar protofilament fold to other *in vitro* aggregated and amplified αSyn, but the global helical organization of the fibrils are distinct.

Finally, the MSA-amplified αSyn fibrils share a similar core as αSyn fibrils directly extracted from brain tissue of MSA patients (Schweighauser et al., 2020). In detail, superimposing the MSA-amplified structure onto the *ex vivo* structure revealed that the triple-stacked *L*-shape is present in both structures (**Figure 3B**). Thus, *in vitro* amplification using brain extracts as seeds and recombinant αSyn as substrate enables replicating αSyn structures found brain lysates of MSA patients partially.

## Discussion

Aggregation of αSyn is the pathological hallmark of both PD and MSA (Goedert, 2015; Goedert, Masuda-Suzukake, & Falcon, 2017). Over recent years, αSyn fibrils prepared in various ways have been suggested as potential toxic species with clinical relevance for PD and MSA (Bousset et al., 2013; Peelaerts et al., 2015). A firm understanding of how αSyn fibrils associated with PD or MSA hamper cellular function, however, has remained elusive. Here we amplified αSyn fibrils from brain extracts of patients pathologically confirmed with PD or MSA, evaluated their potential to seed αSyn-related pathology in oligodendrocytes, and determined their 3D structures by cryo-EM. Our results suggest that MSA-amplified αSyn fibrils are more potent in engendering pathological αSyn assemblies within oligodendroglia than PD-amplified fibrils (**Figure 1**), which is likely the consequence of a different αSyn polymorph.

The 3D-structures of PD- and MSA-amplified αSyn fibrils revealed major differences between both types in their inter-protofilament interfaces, their adopted helical arrangement, and the lacking N-terminal region in MSA-amplified αSyn (**Figure 2**). Interestingly, almost identical αSyn structures were previously solved, namely polymorphs 2A and 2B, originating from *in vitro* aggregated, recombinant αSyn (Guerrero-Ferreira et al., 2019) (**Figure 3**). However, both polymorphs 2A and 2B were observed next to each other and originate from an identical preparation (Guerrero-Ferreira et al., 2019), suggesting that under particular conditions, both polymorphs may denote thermodynamically stable αSyn aggregates. In the present study, we also used recombinant αSyn as the substrate for PMCA. Still, we obtained a single main polymorph in independent and separated experiments, either with seeds from PD or MSA diagnosed brains (**Figure 2**). PMCA is based on the propensity that prions and prion-like proteins act like seeds and replicate in an autocatalytic process, thereby converting recombinant protein as a substrate into amyloid fibrils. Indeed, the PMCA technique was successful in amplifying and detecting misfolded prion proteins implicated in prion diseases (Saa, Castilla, & Soto, 2006; Saborio, Permanne, & Soto, 2001), and several lines of evidence suggest prion-like features also for αSyn (Goedert et al., 2017; Ma, Gao, Wang, & Xie, 2019; Steiner, Quansah, & Brundin, 2018). Considering that the identical and extensively validated PMCA procedure was used to amplify αSyn seeds from either PD or MSA (Strohäker et al., 2019) yields different αSyn structures, suggests that the brain homogenates must contain structurally different αSyn seeds that served as the starting point for amplification leading to the herein described fibrils.

Remarkably, we found that the PD- and MSA-amplified αSyn fibrils have different activity when added to mouse primary oligodendroglial cultures: the MSA-amplified fibrils are more potent in recruiting the endogenous oligodendroglial αSyn and evoking a redistribution of TPPP/p25α protein when compared to the PD-amplified fibrils (**Figure 1**). Both events are characteristic of MSA (Lee, Ricarte, Ortiz, & Lee, 2019). Thus, under physiological conditions, TPPP/p25α is predominant in myelin sheaths (Lehotzky et al., 2010), but under MSA-related pathological conditions, TPPP/p25α relocates to the oligodendrocyte soma (Song et al., 2007). Interestingly, TPPP/p25α not only co-localizes with filamentous αSyn (Kovacs et al., 2004) but, adding to this, also fosters further aggregation of αSyn into filamentous aggregates (Lindersson et al., 2005). The C-terminus of αSyn fibrils has been identified as the binding epitope for TPPP/p25α (Ferreira et al., 2021; Lindersson et al., 2005; Szunyogh, Olah, Szenasi, Szabo, & Ovadi, 2015), but an atomic picture of how TPPP/p25α binds to filamentous αSyn has not been realized yet. Considering the wide range of αSyn fibril polymorphism (**Figure S5**) and that neither in our (**Figure 2**) nor any structurally related fibril (**Figure 3A**; **Table S1, S2**) the C-terminus was resolved, clarification of the full-length αSyn structure under pathological conditions might be necessary to elucidate the underlying structure-activity relationship fully.

Alternatively, the distinct electrostatic surface potentials that discriminate strongly between PD- and MSA-amplified αSyn (**Figure 2E**) also denote a credible source triggering different cellular responses. For example, the reduced electrostatic interactions of the MSA-amplified compared to the PD-amplified αSyn may foster the recruitment of the endogenous oligodendroglial αSyn (**Figure 1**). Such recruitment processes are prerequisites for further fibril elongation, for which hydrophobic interactions of the non-β-amyloid component *NAC* have been identified as essential (Ueda et al., 1993). One might assume that the hydrophobic interactions are weakened under the influence of pronounced electrostatic surface potentials, explaining why our MSA-amplified structure is more potent in endogenous oligodendroglial αSyn recruitment. However, although oligodendrocytes have been shown to internalize αSyn fibrils, oligodendrocytes internalize αSyn monomers and oligomers to a much greater extent (Reyes et al., 2014), suggesting that a direct, intracellular interaction between TPPP/p25α and αSyn fibrils is not necessarily the leading source for promoting TPPP/p25α redistribution. Alternatively, and beyond the scope of this study, one might also consider an indirect effect of αSyn fibrils on TPPP/p25α redistribution and co-localization.

For both amplified fibrils, major differences are found at the N-terminus and in the inter-protofilament interface (**Figure 2**). Assuming that PMCA rigorously amplifies the preformed seeds suggests that the brain homogenates may contain αSyn seeds likely specific for PD or MSA. A fundamental difference between both synucleinopathies is that in the case of PD high concentration of αSyn aggregates is found in dopaminergic neurons, whereas MSA, by contrast, is associated with αSyn inclusions within oligodendrocytes (Alafuzoff & Hartikainen, 2017; Spillantini & Goedert, 2000). Peng *et al*. showed that the intracellular environments of either neurons and oligodendrocytes determine how the same misfolded αSyn seeds develop into different aggregates (Peng et al., 2018). While dopaminergic neurons are directly associated with reward-motivated behavior and motor control, oligodendrocytes are essential for the long-distance saltatory conduction of neuronal impulses, as they enwrap central nervous system axons with the myelin sheath, a lipid enriched multilayer membrane (Montani, 2020; Poitelon, Kopec, & Belin, 2020). Although the lipid content and composition of oligodendrocytes is still unknown, these numbers are well known for the myelin sheath (Norton & Poduslo, 1973; O’Brien, Sampson, & Stern, 1967), which is extremely rich in lipids (∼ 80% of its dry weight) (Montani, 2020; Poitelon et al., 2020). Previous studies revealed the αSyn N-terminus is essential for lipid-binding (Davidson et al., 1998) and fatty acid-induced oligomerization (Karube et al., 2008). Further biophysical experiments suggest that αSyn-lipid interactions are predominantly driven by electrostatic interactions between charged N-terminal residues and the charged head groups of phospholipids (Jo, McLaurin, Yip, St George-Hyslop, & Fraser, 2000), which comprise ∼26% of all lipids found in central nervous system myelin (Norton & Poduslo, 1973). Thus, one might assume that the αSyn N-terminus interacts with these phospholipids during fibril aggregation and seed formation in oligodendrocytes, which, in turn, may hamper the stabilization of the folded N-terminus in these cells. Indeed, previous site-directed spin labeling and electron paramagnetic resonance spectroscopy also revealed a structurally heterogeneous N-terminus (aa 1 - 30) in αSyn fibrils (Der-Sarkissian et al., 2003). Although no lipids were present during *in vitro* amplification (Strohäker et al., 2019), PMCA rigorously replicates the visible seeds, in which the N-terminus was apparently not properly stabilized. This assumption is further corroborated by our local resolution estimates. The reconstructed densities of the MSA-amplified αSyn suggest a high local resolution at the inter-protofilament interface. Still, the estimated resolution gets lower towards the periphery of the fibril (**Figure S3**), meaning a higher degree of structural flexibility in these particular regions.

Although it is reasonable to assume that PMCA rigorously amplifies the αSyn seeds, a recent study suggests that this is not necessarily the case (Lövestam et al., 2021). Initially, the authors determined the atomic structures of αSyn fibrils extracted from the putamen of MSA-diagnosed patients (Schweighauser et al., 2020). These preparations were subsequently used for *in vitro* seeded assembly of recombinant αSyn using PMCA. While the obtained structures are similar to our MSA-amplified structure, they are at first glance quite different from those of the *ex vivo* seeds (Lövestam et al., 2021). However, it is remarkable that the amplified and *ex vivo* protofilament structures can in fact be partially aligned in an anti-parallel arrangement at least within a triple-stacked *L*-shape core region comprising about 40 residues (**Figure 3B**). Even though the *ex vivo* structures are asymmetrical (Schweighauser et al., 2020) (**Figure S6**), which might be due to additional co-solutes bound in their protofilament interface, it is conceivable that this matching 40 residue core region acts as the dominant seed and is therefore replicated in the amplified structures.(Lövestam et al., 2021; Schweighauser et al., 2020)

It should be noted that any discrepancies between our amplified fibril structures and those determined by Lövestam et al. (Lövestam et al., 2021) could be due to the fact that we extracted αSyn from the human amygdala (Strohäker et al., 2019), while Schweighauser *et al*. extracted αSyn from putamen (Schweighauser et al., 2020). Furthermore, considering that several different folds for αSyn fibrils are known (**Figure S5, Table S3**), one might speculate that different brain regions might accumulate different dominant αSyn polymorphs. Still, additional analyses are required to corroborate this assumption more firmly. In sum, *in vitro* amplification using brain extracts as seeds and recombinant αSyn as substrate enables partial replication of αSyn assemblies found brain lysates of MSA patients.

Taken all together, we report two αSyn structures extracted and amplified from brain tissue of PD- and MSA-confirmed individuals. Both types of αSyn fibrils share the same proto-filament fold and reveal differences in their inter-protofilament interfaces, their adopted helical arrangement, and at the N-terminal region, allowing discrimination between PD- and MSA-amplified αSyn fibrils. Furthermore, the MSA-amplified αSyn fibrils bear an increased potency to seed αSyn-related pathology and to evoke p25α re-distribution, events underlying oligodendroglial dysfunction. Thus, our observations are consistent with the one disease-one strain hypothesis, assuming a unique connection between the clinical disease presentation and a single, defined structure of aSyn fibril structure. However, it is out of the question that more *in vitro* amplified and *ex vivo* αSyn structures must be determined at high resolution. Still, it is equally essential to investigate the functional implications of those fibrils in more detail. The combination of functional data with structural insights as pursued in the current study bears enormous potential and opens up a new avenue to understand the sophisticated pathologic mechanisms of α-synucleinopathies.

## Materials & Methods

### Amplification of *α*Syn fibrils from PMCA-products

Ethics approval for the study of brain tissue was from the University of New South Wales Human Research Ethics Committee (approval number: HC16568). αSyn aggregates used in the current study had been previously amplified from brain extracts of patients pathologically confirmed with PD and MSA using PMCA (Strohäker et al., 2019). To obtain sufficient quantities for structural analysis, PMCA-amplified amyloid fibrils were used in a second step to seed recombinant αSyn (Strohäker et al., 2019).

N-terminally acetylated αSyn was obtained by co-transfection of *E. coli* BL21 (DE3) cells with pT7-7 plasmid encoding for human αSyn (kindly provided by the Lansbury Laboratory, Harvard Medical School, Cambridge, MA) and *S. pombe* NatB acetylase complex (Johnson, Coulton, Geeves, & Mulvihill, 2010) using pNatB plasmid (pACYCduet-naa20-naa25, Addgene, #53613, kindly provided by Dan Mulvihill). Protein expression and purification was performed as described (Hoyer et al., 2002).

αSyn fibrils were prepared at 37 °C from monomeric, N-terminally acetylated αSyn taken from the supernatant of freshly thawed αSyn on ice after ultracentrifugation (Beckman Coulter Optima MAX-XP using a TLA 100.3 rotor with a rotor speed of 55,000 rpm). 0.5% (w/w) PMCA product was added to 250 μM αSyn stock solution (50 mM HEPES, 100 mM NaCl, pH 7.4, 0.02% NaN_3_) and initially water bath sonicated for 10 minutes. This mixture was aggregated under quiescent conditions in 1.5 mL Eppendorf cups in a ThermoScientific Heratherm incubator. The αSyn fibrils imaged by cryo-EM were taken from the sample that was applied to primary oligodendroglial cultures.

### Primary oligodendroglial cultures

Mixed glial cultures generated from P0 to P3 neonatal wild-type (WT) mice were maintained in full DMEM for 10 to 14 days until a monolayer of astrocytes on the bottom and primary oligodendroglial progenitor cells (OPCs) with loosely attached microglia on the top, were apparent. The separation of OPCs was achieved initially with the removal of microglia, by shaking in 200 rpm for 1h in 37°C and then with continuous shaking under the same conditions for 18 hrs, as previously described (Mavroeidi et al., 2019). Afterwards, isolated cells were platted on poly-D-lysine-coated coverslips (P7405, Sigma-Aldrich, USA) with a density of 80,000 cells/mm2 and maintained in SATO medium (284369) supplemented with Insulin-Transferrin-Selenium solution (41400045, ITS-Gibco, Invitrogen, Carlsbad, CA, USA), 1% penicillin/streptomycin and 1% horse serum (H1138; Sigma-Aldrich, St. Louis, MO, USA) for 4 days. αSyn fibrils (final concentration 0.5 µg/mL culture medium/well) amplified from human MSA and PD brains were added to TPPP/p25α-positive mature differentiated oligodendrocytes for 48 hrs and then cells were fixed and preceded for immunofluorescence analysis.

### Immunocytochemistry and confocal microscopy

Forty-eight (48) hours post patient-amplified fibril addition, cells were fixed with 4% paraformaldehyde for 40 min, blocked in 10% normal goat serum containing 0.4% Triton X-100 for 1 h at room temperature, and incubated with antibodies against the human (LB509), the rodent (D37A6), or the oxidized/nitrated (Syn303) αSyn and the oligodendroglial phospho-protein TPPP/p25α (kind gift from Dr. Poul Henning Jensen, Aarhus University, Denmark) overnight at 4 °C. Images were obtained using a Leica TCS SP5 confocal microscope combined with a dual (tandem) scanner. All confocal images were obtained under equivalent conditions of laser power, pinhole size, gain, and offset settings between the groups. ImageJ (v2.0.0) software was used to quantify relative protein levels expressed as % area coverage, normalized to the p25α+ cells/field.

### Cryo-EM grid preparation and imaging

Sample volumes of 3.5 µl were applied to freshly glow-discharged R3.5/1 holey carbon grids (Quantifoil) and vitrified using a Mark IV Vitrobot (Thermo Fischer Scientific) operated at 100% rH and 20 °C. Micrographs were collected with a Titan Krios transmission-electron microscope operated at 300 keV accelerating voltage at a nominal magnification of 81,000 x using a K3 direct electron detector (Gatan) in non-superresolution counting mode, corresponding to a calibrated pixel size of 1.05 Å on the specimen level. In total, 6,780 images (as to PD associated fibrils) and 4,474 (as to MSA relevant fibrils) with defocus values in the range of - 0.7 µm to -2.9 µm were recorded in movie mode with 2.2 s acquisition time. Each movie contained 40 frames with an accumulated dose of approximately 43 electrons per Å^2^. The resulting dose-fractionated image stacks, containing all frames 1-40, were subjected to beam-induced motion correction using MotionCor2 (Zheng et al., 2017), prior to helical reconstruction. Estimation of contrast transfer function parameters for each micrograph was performed using CTFFIND4 (Rohou & Grigorieff, 2015). Subsequently, PD and MSA related αSyn fibrils were reconstructed using RELION-3.1 (Zivanov, Nakane, & Scheres, 2020), following the helical reconstruction scheme (He & Scheres, 2017).

### Helical reconstruction of PD-amplified fibrils

crYOLO (Wagner et al., 2019) was used for the selection of 76,170 fibrils in the dataset, from which 2,223,091 segments were extracted using a 19 Å inter-box distance. Using RELION-3.1 (Zivanov et al., 2020), maximum-likelihood two-dimensional (2D) reference-free classification, and 3D classification were performed on an unbinned data set (1.05 Å/px, 200px box size); a cylinder with white noise added using EMAN2 (Tang et al., 2007) was used for the initial reference.

We used the CHEP algorithm (Pothula, Geraets, Ferber, & Schröder, 2021; Pothula, Smyrnova, & Schröder, 2019) with *k*-means clustering on the results of 2D classification to identify and group fibrils by overall conformation (*k* = 3), and investigate these 3 clusters individually. The algorithm associates segments extracted from a single helical fibril and clusters each fibril together with similar fibrils, based on the similarity of their segments within the classification. On a single-subunit level the signal-to-noise ratio is often not enough to distinguish between two similar classes, especially for some projections where the conformations appear more similar.

After iterative classification steps, 23,937 particles were selected for 3D auto-refinement, beam tilt refinement, CTF refinement, and reconstruction in RELION-3.1 (Zivanov et al., 2020). During the postprocessing step in RELION, the map was masked with a soft mask and a B-factor of −112 Å^2^ was applied, and the resolution was estimated as 3.3 Å based on the gold-standard Fourier shell correlation 0.143 criterion (**Figure S2A**). The helical geometry was then applied to the map, which was then re-sharpened using VISDEM (Spiegel, Duraisamy, & Schröder, 2015). Local resolution was determined using RELION-3.1 (Zivanov et al., 2020).

### Helical reconstruction of MSA-amplified fibrils

crYOLO (Wagner et al., 2019) was used for the selection of 101,588 fibrils in the dataset, from which 1,848,677 segments were extracted using a 50 Å inter-box distance and RELION-3.1 (Zivanov et al., 2020) was used for reconstruction. To exclude non-twisted and irregularly twisted segments, we initially performed several rounds of 2D classification on a downscaled data set (2.1 Å/pix, 500 pix box size). For 3D classification we (re-)extracted 91,552 segments without downscaling (1.05 Å/pix, 250 pix box size).

Starting from a featureless cylinder filtered to 60 Å, we performed a 3D classification with one class and *T* = 3, followed by a 3D classification with one class and *T* = 4, after which the two protofilaments were separated and first backbone features became visible. Subsequently, another round of 3D classification with five classes and *T* = 4 yielded one class showing clear backbone features. This class was selected (25,692 particle segments) for further rounds of 3D classification (*k* = 1) with step-wise adjusting the *T*-value from 4 to 8 after which separation of β-strands along the Z-axis and the approximate 2_1_ screw symmetry between the two protofilaments became visible. From here on, we performed multiple rounds of 3D auto-refinement until no further improvement of the map was observed. Assuming a left-handed twist, the helical twist and rise converged to 179.66° and 2.37 Å, respectively, in agreement with the predominant crossover distances measured on the motion-corrected cryo-EM micrographs. Finally, post-processing with a soft-edged mask and an estimated sharpening *B*-factor of -101 Å^2^ yielded post-processed maps. The resolution was estimated from the value of the FSC curve for two independently refined half-maps at 0.143 (**Figure S2B**). The helical geometry was then applied to the map, which was then re-sharpened using VISDEM (Spiegel et al., 2015). Local resolution was determined using RELION-3.1 (Zivanov et al., 2020).

### Atomic model building and refinement

PDB entries 6SSX and 6SST (Guerrero-Ferreira et al., 2019) were used for an initial model towards αSyn extracted from PD and MSA brain, respectively. Subsequent refinement in real space was conducted using PHENIX (Afonine et al., 2018; Liebschner et al., 2019). With PD, the final refined protofilament subunit had an RMSD of 0.64 Å to PDB 6SSX. As for MSA, the final refined protofilament subunit had an RMSD of 0.95 Å to PDB 6SST.

### Determination of electrostatics for *α*Syn fibrils

We calculated the electrostatics for αSyn fibrils using the APBS/PDB2PQR server (https://server.poissonboltzmann.org/) (Baker, Sept, Joseph, Holst, & McCammon, 2001; Dolinsky, Nielsen, McCammon, & Baker, 2004). Therefore, we used our atomic models and assembled 60 peptides into a fibril, imposing the helical symmetry reported in **Table 1**. For interpretation of the results, we focused on the central slices of the fibrils. APBS calculations for polymorphs 2A and 2B (Guerrero-Ferreira et al., 2019) were conducted analogously.

## Acknowledgements

B.F., J.A.G., and G.F.S. are grateful for computational support and infrastructure provided by the ‘‘Zentrum für Informations-und Medientechnologie’’ (ZIM) at the Heinrich Heine University Düsseldorf and the computing time provided by Forschungszentrum Jülich on the supercomputer JURECA/JURECA-DC at Jülich Supercomputing Centre (JSC). Brain tissues (approval number: PID399) were received from the Sydney Brain Bank at Neuroscience Research Australia which is supported by The University of New South Wales, Neuroscience Research Australia and Schizophrenia Research Institute. M.Z. was supported by the European Research Council (ERC) under the EU Horizon 2020 research and innovation programme (grant agreement No. 787679). Access to the FEI Titan Krios cryo-electron microscope with K3 detector was gratefully provided by Prof. Dr. Patrick Cramer at the MPI for Biophysical Chemistry, Goettingen, Germany.

## Authors contributions

M.Z. and G.F.S conceived the study; W.S.K provided the brain samples; B.C.J and S.-J.L. amplified aSyn seeds from brain tissue using PMCA; T.S. conducted protein preparation, and fibril amplification; P.M. and M.X. performed the cell experiments, and analyzed the associated data; C.D. conducted cryo-EM grid preparation and data collection; B.F., J.A.G., and G.F.S. performed image processing, reconstruction and model building; B.F., J.A.G, M.Z., and G.F.S. wrote the paper. All authors discussed results and commented on the paper.

