## Supplemental Material for "α-Synuclein polymorphism determines oligodendroglial dysfunction"

### Supplemental Figures

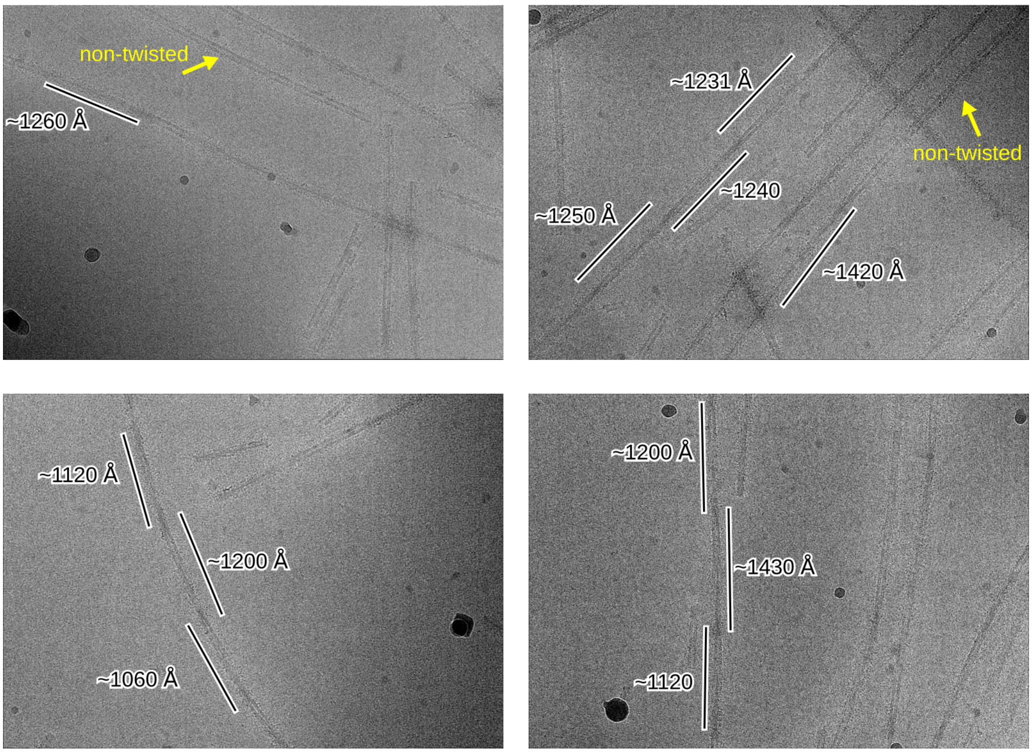


**Figure S1: Representative micrographs with MSA-type αSyn fibrils.**
Micrographs showing twisted (labeled with cross over distances) and non-twisted αSyn fibrils amplified from MSA patient's material.

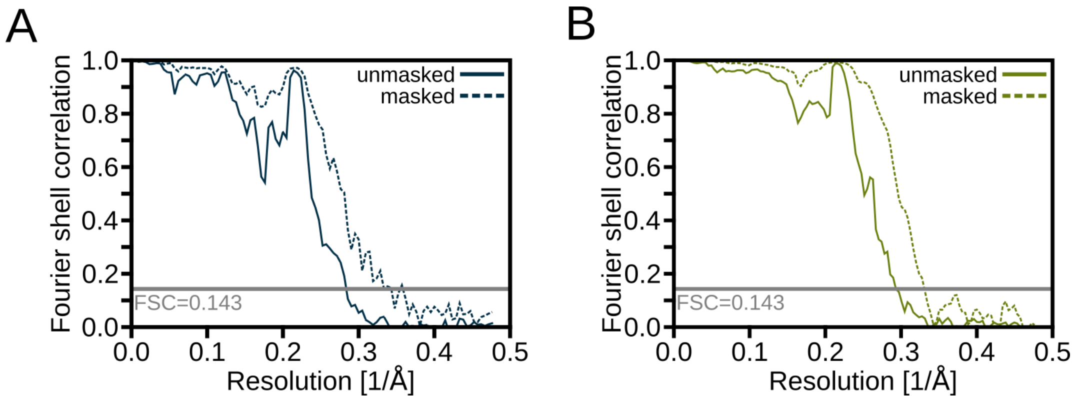


**Figure S2: Fourier shell correlation curves.**

The Fourier shell correlation curves are shown between two independently refined unmasked (solid lines) and masked (dashed lines) half-maps. The *z*-percentage was 0.3 in the case of PD (**A**) and 0.1 in the case of in MSA (**B**), respectively.

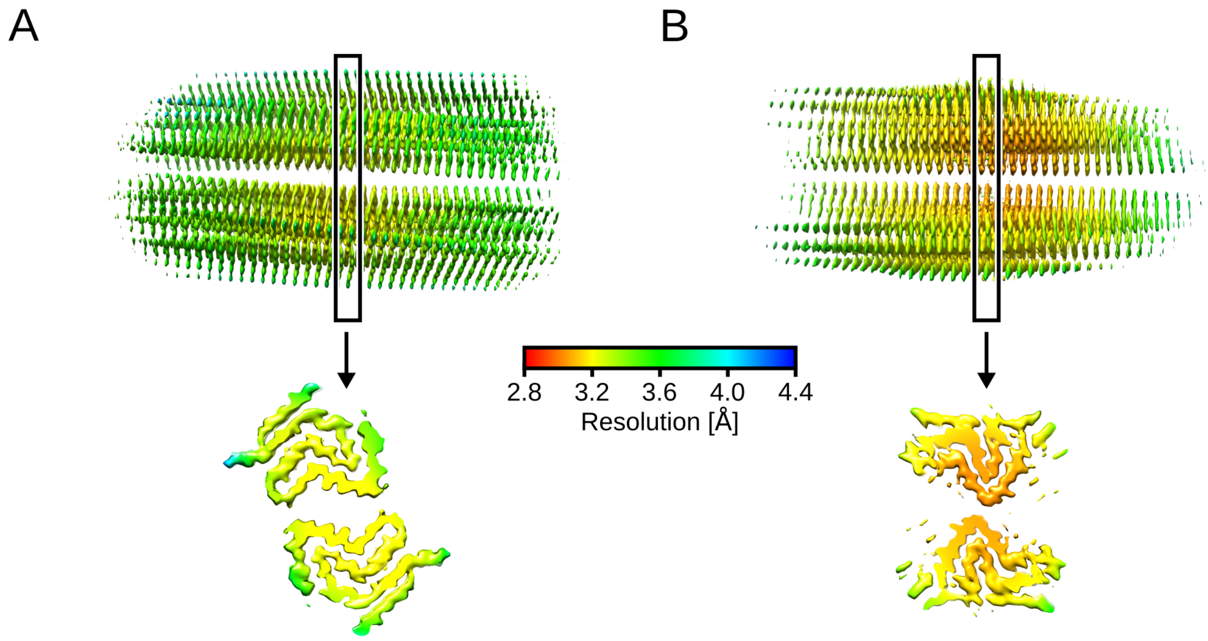


**Figure S3: Local resolution estimation.**
Reconstructed map of PD (**A**) and MSA (**B**) fibrils colored according to the local resolution estimation (see color scale). The lower panel shows a cross-section of the central region.

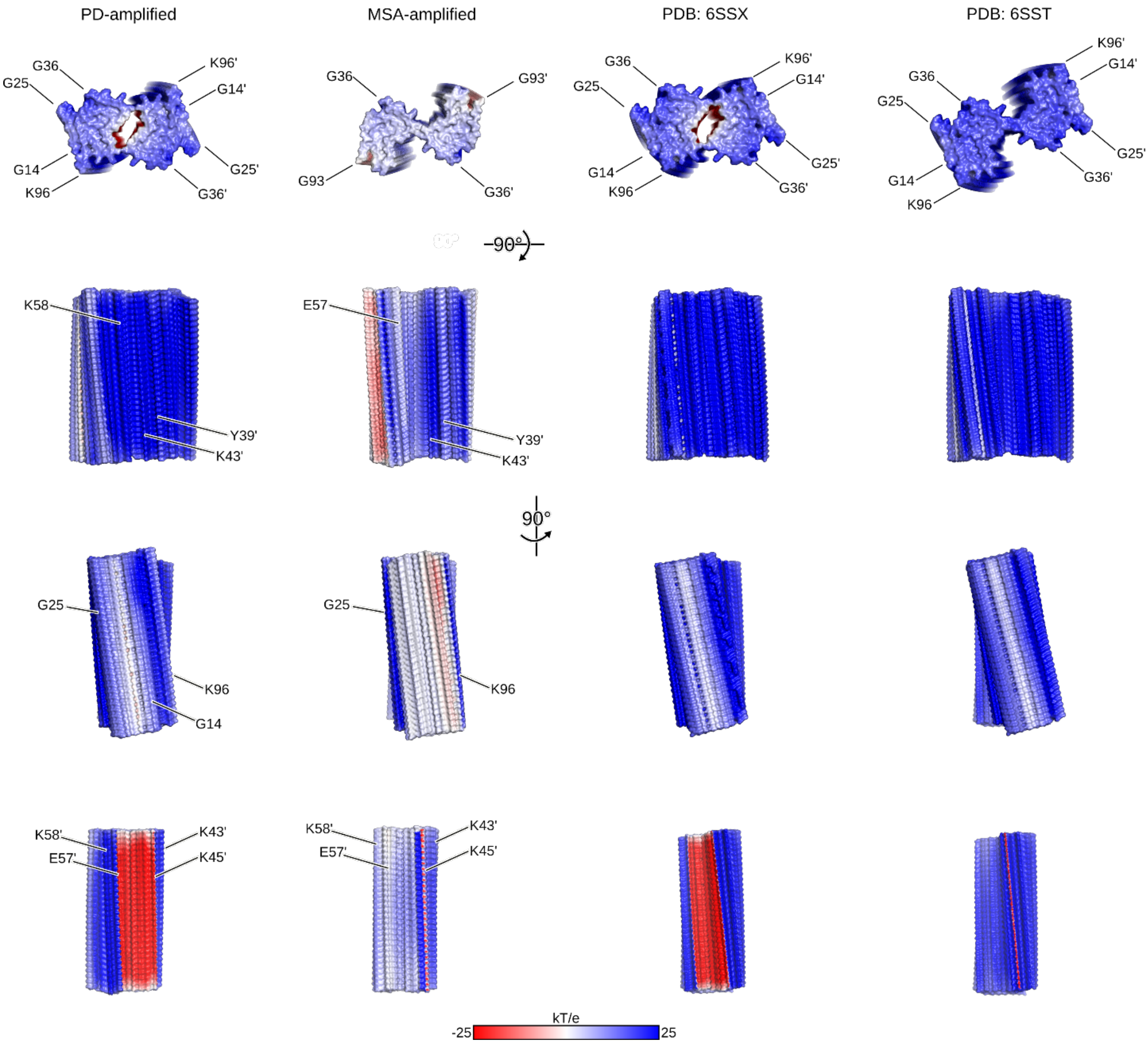


**Figure S4: Electrostatic surface potential.**
The cryo-EM structures of amplified αSyn fibrils from PD (left) and MSA (right) brain and two αSyn fibrils of in vitro aggregated recombinant protein (Guerrero-Ferreira et al., 2019) viewed from different angles. The calculated electrostatic potential was mapped onto the surface and colored according to the color scale at the bottom. The bottom panel shows a cross section along the Z-axis, with the inter-protofilament interfaces now oriented towards the viewer. For illustrative purposes, some amino acids are explicitly labeled, with amino acids from subunit B marked with an additional prime.

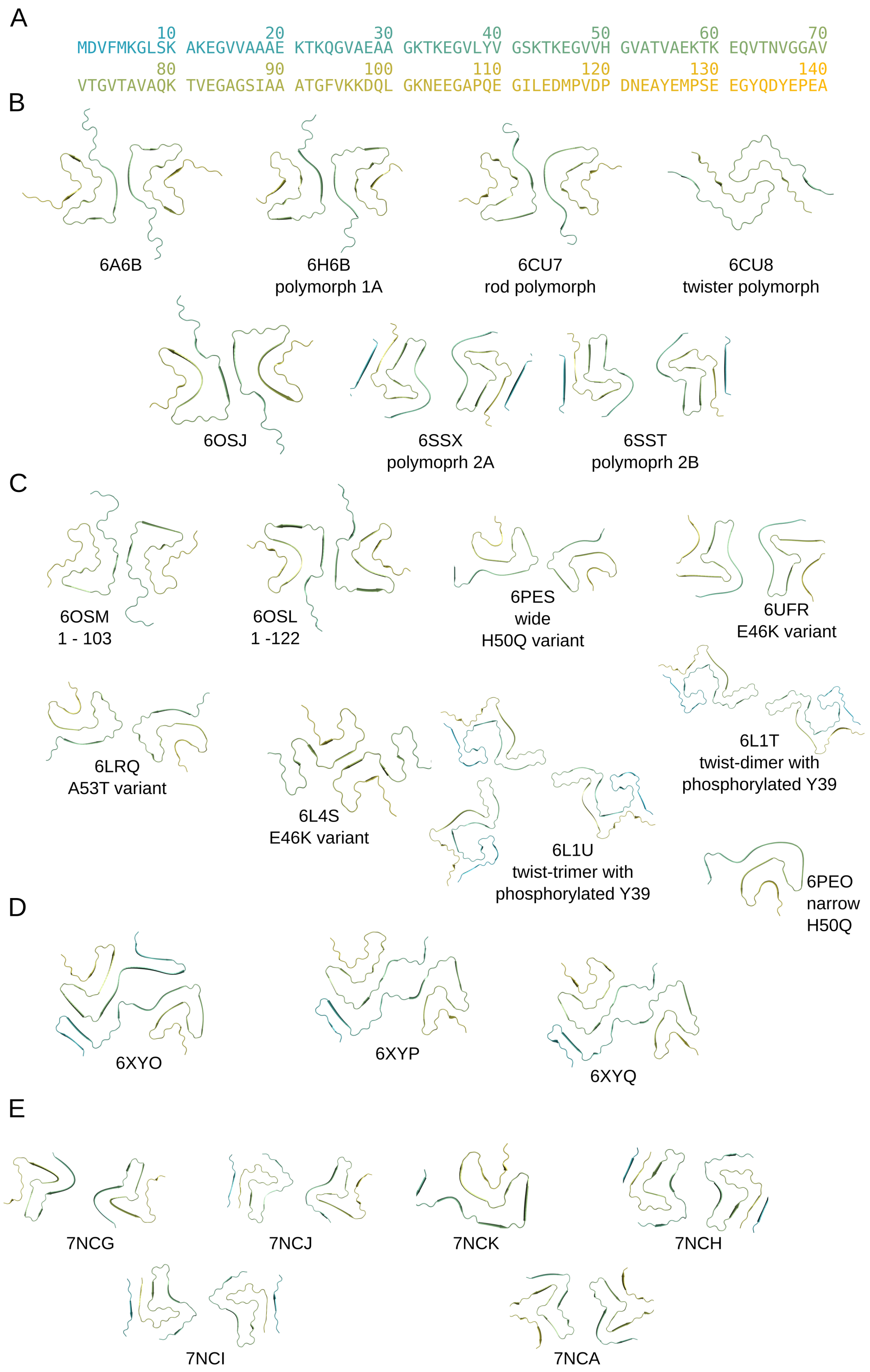
A
10 20 30 40 50 60 70
MDVFMKGLSK AKEGVVAAAE KTKQGVAEAA GKTKEGVLYV GSKTKEGVVH GVATVAEKTK EQVTNVGGAV
80 90 100 110 120 130 140
VTGVTAVAQK TVEGAGSIAA ATGFVKKDQL GKNEEGAPQE GILEDMPVDP DNEAYEMPSE EGYQDYEPEA
B
6A6B
6H6B
polymorph 1A
6CU7
rod polymorph
6CU8
twister polymorph
6OSJ
6SSX
polymoprh 2A
6SST
polymoprh 2B
C
6OSM
1 - 103
6OSL
1 -122
6PES
wide
H50Q variant
6UFR
E46K variant
6LRQ
A53T variant
6L4S
E46K variant
6L1T
twist-dimer with
phosphorylated Y39
6L1U
twist-trimer with
phosphorylated Y39
6PEO
narrow
H50Q
D
6XYO
6XYP
6XYQ
E
7NCG
7NCJ
7NCK
7NCH
7NCI
7NCA

**Figure S5: Summary of previously resolved αSyn structures.**

**A:** Full-length mino acid sequence of human αSyn (UniProt: P37840). The sequence is colored from the N- to the C-terminus according to the blue-green-yellow pallet. **B-E:** Top view onto the previously resolved αSyn structures. The structures are colored according to the color code in **A**. The four-letter PDB-ID is reported with the structures. The structures can be organized into four groups, in which the fibrils are formed by recombinant full-length wild type aSyn (**B**), recombinant, truncated, or modified aSyn (**C**), aSyn extracted from brain tissue of MSA diagnosed patients (**D**), and aSyn amplified from seeded brain extracts (**E**) (for details, please see **Table S3**)

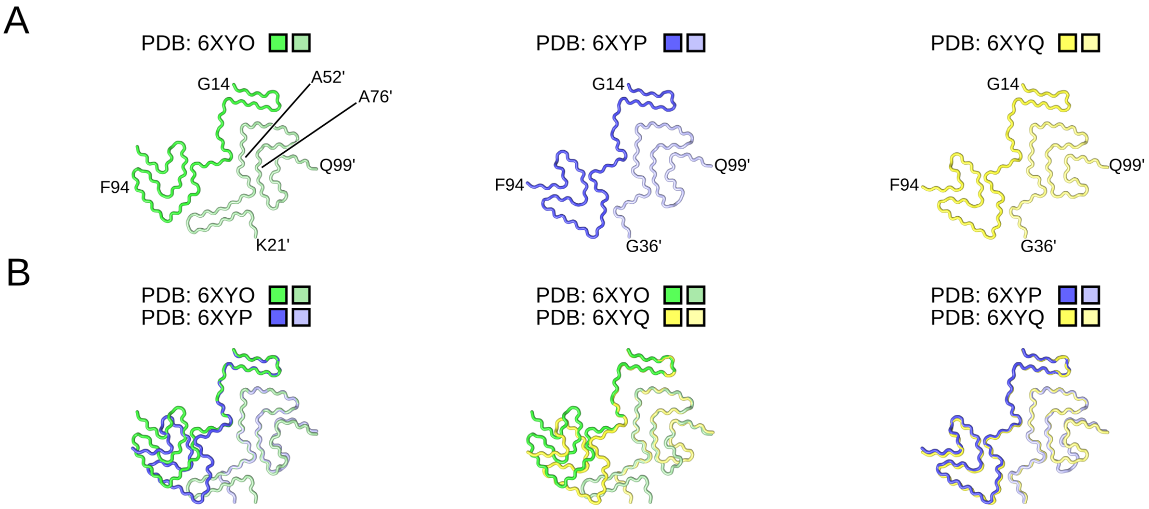


**Figure S6: Comparison between *ex vivo* MSA type αSyn fibrils.**
**A**: Top view onto the αSyn structures from the brains of individuals with MSA (Schweighauser et al., 2020). The protofilaments are colored in different shades of green, blue, and yellow, with amino acids from subunit B labeled with a prime. **B**: Superposition of αSyn structures from **A.** All structures were superimposed on the region extending from A52' to A57'.

#### Supplemental Tables

**Table S1. PD-amplified αSyn fibril structure compared to the *in vitro* polymorph 2A (PDB ID 6SSX (Guerrero-Ferreira et al., 2019)) and MSA-amplified fibril structure from Lovestam et al. (PDB ID 7NCH (Lövestam et al., 2021)).**

| | **PD-amplified αSyn** | **PDB 6SSX** | **PDB 7NCH** |
|---|---|---|---|
| Overlay onto PD-amplified αSyn[a] | 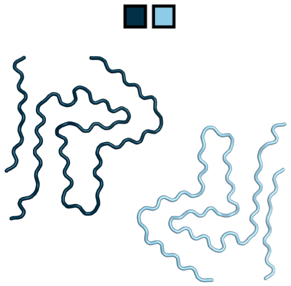 | 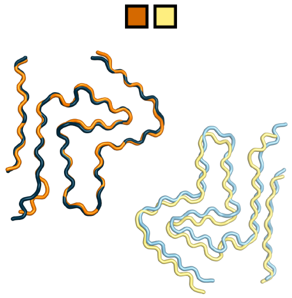 | 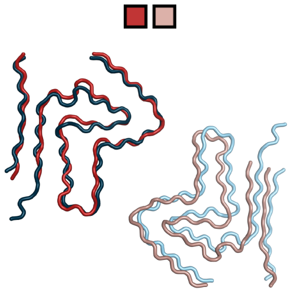 |
| Cα RMSD to PD type αSyn [Å] (one/two protofilament)[b] | -- / -- | 1.03 / 1.33 | 1.65 / 2.65 |
| Symmetry | C2 | C2 | C2 |
| Rise [Å] / Twist [°] | 4.68 / -0.78 | 4.80 / -0.80 | 4.78 / -0.86 |

[a] To visualize the displacement between two opposite protofilaments, only the Cα atoms of one protofilament (shown in darker colors) were aligned.
[b] Only amino acids present in both structures (mobile and reference) were considered for RMSD calculations.

**Table S2. MSA-amplified αSyn fibril structure compared to the *in vitro* polymorph 2B (PDB ID 6SST (Guerrero-Ferreira et al., 2019)) and MSA-amplified fibril structure from Lovestam et al. (PDB IDs 7NCI and 7NCG (Lövestam et al., 2021)).**

| | **MSA type αSyn** | **PDB 6SST** | **PDB 7NCI** | **PDB 7NCG** |
|---|---|---|---|---|
| Overlay onto MSA type αSyn[a] | 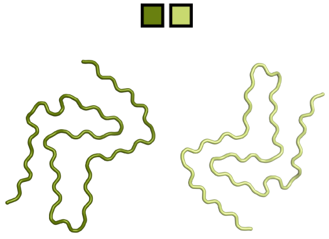 | 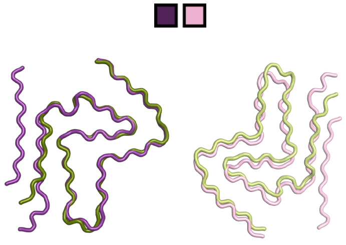 | 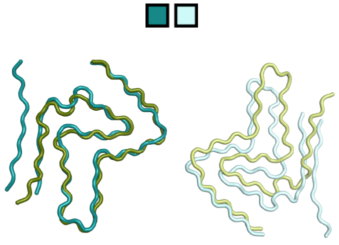 | 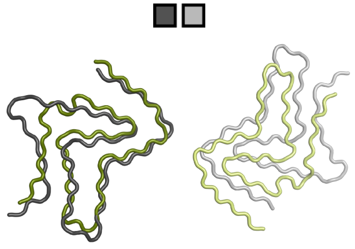 |
| Cα RMSD to MSA type αSyn [Å] (one/two protofilament)[b] | -- / -- | 1.48 / 1.74 | 1.50 / 12.55 | 4.57 / 9.23 |
| Symmetry | C1 | C1 | C2 | C2 |
| Rise [Å] / Twist [°] | 2.37 / 179.66 | 2.40 / 179.55 | 4.75 / -0.77 | 4.75 / -0.95 |

[a] To visualize the displacement between two opposite protofilaments, only the Cα atoms of one protofilament (shown in darker colors) were aligned.
[b] Only amino acids present in both structures (mobile and reference) were considered for RMSD calculations.

**Table S3: Summary of αSyn structures.**

| PDB-ID | Polymorph name | Variation | Ref. |
|---|---|---|---|
| **Recombinant full-length human αSyn** | | | |
| 6FLT (see 6H6B) | | full-length human αSyn | |
| 6A6B | | full-length human αSyn, | (Y. Li et al., 2018) |
| 6H6B | polymorph 1A | full-length human αSyn | (Guerrero-Ferreira et al., 2018) |
| 6CU8 | Twister/ poly-morph 1B | full-length human αSyn | (B. Li et al., 2018) |
| 6CU7 | Rod | full-length human αSyn | (B. Li et al., 2018) |
| 6RTB (see 6SST ) | polymorph 2B | full-length human αSyn | |
| 6RT0 (see 6SSX) | polymorph 2A | full-length human αSyn | |
| 6OSJ | | full-length human αSyn | (Ni, McGlinchey, Jiang, & Lee, 2019) |
| 6SSX | polymorph 2A | full-length human αSyn | (Guerrero-Ferreira et al., 2019) |
| 6SST | polymorph 2B | full-length human αSyn | (Guerrero-Ferreira et al., 2019) |
| **Recombinant truncated or modified aSyn** | | | |
| 6OSM | | Recombinant 1 -103 human αSyn | (Ni et al., 2019) |
| 6OSL | | Recombinant 1 - 122 human αSyn | (Ni et al., 2019) |
| 6PES | wide | Recombinant full-length H50Q human αSyn | (Boyer et al., 2019) |
| 6PEO | narrow | Recombinant full-length H50Q human αSyn | (Boyer et al., 2019) |
| 6UFR | | Recombinant full-length E46K human αSyn | (Boyer et al., 2020) |
| 6LRQ | | Recombinant full-length A53T human αSyn | (Sun et al., 2020) |
| 6L4S | | Recombinant full-length E46K human αSyn | (Zhao, Li, et al., 2020) |
| 6L1T | twist-dimer fibril | Recombinant full-length phosphorylated Y39 human αSyn | (Zhao, Lim, et al., 2020) |
| 6L1U | twist-trimer fibril | Recombinant full-length phosphorylated Y39 human αSyn | (Zhao, Lim, et al., 2020) |
| **aSyn extracted from brain tissue of MSA diagnosed patients** | | | |
| 6XYO | MSA type II-1 | MSA brain extracts, *ex vivo* | (Schweighauser et al., 2020) |
| 6XYP | MSA type I | MSA brain extracts, *ex vivo* | (Schweighauser et al., 2020) |
| 6XYQ | MSA type II-2 | MSA brain extracts, *ex vivo* | (Schweighauser et al., 2020) |

**aSyn amplified from seeded brain extracts**

| | | | |
|---|---|---|---|
| 7NCG | Type 2A | Full-length alpha-synuclein fibril seeded *in vitro* by fibrils purified from MSA brain | (Lövestam et al., 2021) |
| 7NCJ | Type 2AB | Full-length alpha-synuclein fibril seeded *in vitro* by fibrils purified from MSA brain | (Lövestam et al., 2021) |
| 7NCK | Type 3 | Full-length alpha-synuclein fibril seeded *in vitro* by fibrils purified from MSA brain | (Lövestam et al., 2021) |
| 7NCH | Type 1B | Full-length alpha-synuclein fibril seeded *in vitro* by fibrils purified from MSA brain | (Lövestam et al., 2021) |
| 7NCI | Type 2B | Full-length alpha-synuclein fibril seeded *in vitro* by fibrils purified from MSA brain | (Lövestam et al., 2021) |
| 7NCA | Type 1A | Full-length alpha-synuclein fibril seeded *in vitro* by fibrils purified from MSA brain | (Lövestam et al., 2021) |
